# Multi-agentic system for primer design in qPCR and LAMP diagnostics tests

**DOI:** 10.64898/2026.09.15.751771

**Authors:** Kenny J.X. Lau

## Abstract

Primer design is a fundamental component of molecular diagnostics in both quantitative polymerase chain reaction (qPCR) and loop-mediated isothermal amplification (LAMP) assays. However, assay design is often performed manually as nucleotide databases, sequence alignment tools and resources are found at different places on the Internet. In this study, an AI-orchestrated bioinformatics workflow was developed to automate the end-to-end qPCR and LAMP primers and probes. The workflow was implemented using LangGraph, LangChain and Biopython, where a series of specialized agents were coordinated to execute sequential bioinformatics tasks with minimal human intervention.

Target sequences were then retrieved based on the user’s request from the National Center for Biotechnology Information nucleotide database and the requested sequence records were then subjected to multiple sequence alignment for the identification of conserved genomic regions. The multi-agentic primer design system can be used for assay development for applications in infectious disease diagnostics, outbreak surveillance and environmental monitoring. This study also demonstrates how multi-agentic systems can be combined with established bioinformatics methods to automate qPCR and LAMP assay design.

## INTRODUCTION

Quantitative real-time PCR (qPCR) and loop-mediated isothermal amplification (LAMP) are well-established methods for the quantification and species identification of bacteria and fungi in microbiology [1, 2]. Nucleic acid amplification tests (NAATs) have become essential across the life sciences and are now regarded as the gold standard for species detection from a wide range of applications from clinical use [3], food contamination [4] to environmental monitoring [5].

Target specificity is one of the most critical characteristics of a pair of PCR primers and probe in qPCR and 6 primer sets targeting 8 regions in LAMP assays. An ideal primer set should only selectively amplify the intended target sequence while avoiding amplification of any non-target regions [6]. This requirement is particularly important in both assays where accurate target detection depends on highly specific amplification. In SYBR Green assays, fluorescence is generated when the dye binds to double-stranded DNA where the signal detected cannot be distinguished between specific and non-specific amplification [7]. In TaqMan assays, fluorescence is generated when DNA polymerase cleaves the probe during extension, separating the reporter dye from the quencher [8]. Similarly, LAMP assays also employ fluorescent dyes or probes to monitor amplification in real time [9]. Thus, any non-specific amplification can contribute to overall signal that may lead to false positive results, reduced assay specificity and inaccurate target detection.

There are many online tools available for primer design, but the workflow remains fragmented. Users typically need to access sequence databases, download target sequences, perform sequence alignments and identify consensus regions before initiating primer design. Some software tools also require commercial license and uploading of proprietary sequence data to external servers like PrimerPooler, Phuser and Benchling [10-12]. Primer3 is one of the most widely adopted platforms as it is based on a comprehensive set of user-defined parameters including primer length, melting temperature, GC content, and product size [13]. However, Primer3 does not assess target specificity during primer design, requiring users to perform additional analyses using separate tools to evaluate off-target amplification. This multi-step workflow can be laborious when they are large number of candidate primers from sequence matches in database searches. To address the limitations of Primer3, several tools such as *in-silico* PCR and reverse ePCR have been developed to predict amplification targets of user-supplied primer pairs [14, 15]. However, these tools are designed for conventional PCR and are generally not applicable to LAMP assays, which require multiple primers targeting several regions of the template. Consequently, LAMP primer design and specificity assessment often require dedicated software such as NEB LAMP Primer Design Tool [16] and PrimerExplorer [17].

## RESULTS

### Proposed agentic AI framework for primer design

A hierarchical multi-agent framework with two branches, qPCR and LAMP pathways as shown in **Fig 1** highlights a procedural step-by-step workflow of coordinated activities of seven specialized sub-agents. The process starts with a user request where a person may ask for TaqMan and/or LAMP assays for a particular gene in which organism. This could be bacteria, fungi or even viruses if there are matches against the sequence records existing in NCBI nucleotide database. Due to the free-tier limit when hosting on a ZeroGPU space on Huggingface, a smaller size LLM such as the Qwen2.5-1.5 billion parameters trained LLM was used to extract keywords from the user’s request and parse into a JSON string as follows: {{“organism”: “Bacillus”, “gene”: “16S rRNA”, “target_kw”: “subtilis”, “min_amp”: 100, “max_amp”: 300, “min_tm”: 55, “min_probe_tm”: 60}}.

**Fig 1.**
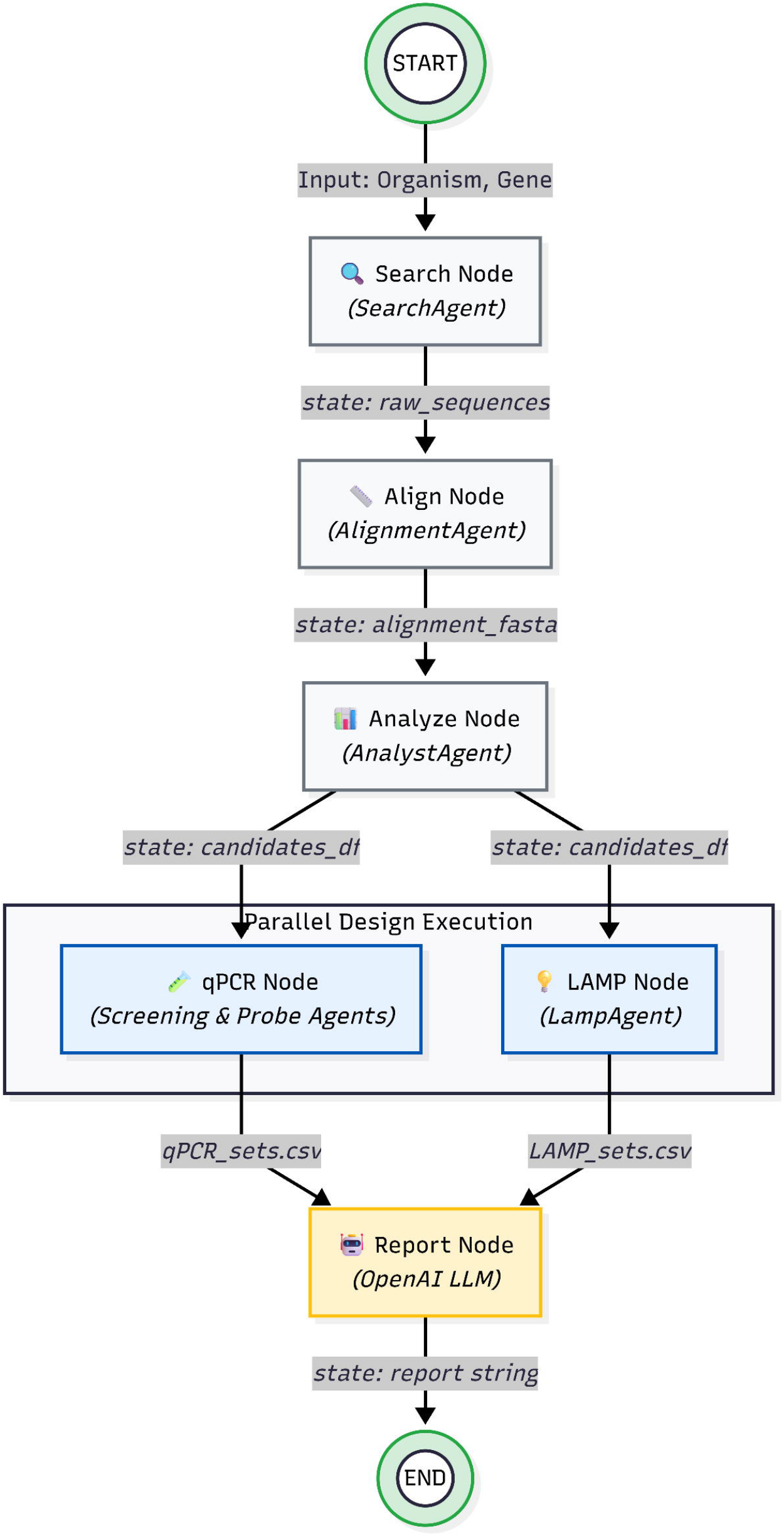
Workflow of multi-agent primer design tool. Input organism and gene information are used to retrieve sequences (SearchAgent), perform sequence alignment (AlignmentAgent) and identify candidate target regions (AnalystAgent). Candidate regions are then processed in parallel for qPCR and LAMP primer design. The outcome and primer set shortlisted will be explained and reported by an LLM.

### Optimising search results from NCBI in sub-agent 1: Search Agent

Querying the NCBI database and retrieving the correct sequence records require the right keywords and parameters. The input is optimised for 16S or Internal Transcribed Spacer (ITS) gene searches, where various user inputs are considered and will be standardized in lower case letters as follows:

if “16s” in gene_lower:

> gene_query = ‘(“16S ribosomal RNA”[Title] OR “16S rRNA”[Title] OR “16S”[Gene])’
>
> elif “its” in gene_lower or “internal transcribed spacer” in gene_lower:
>
> gene_query = ‘(“internal transcribed spacer”[Title] OR “ITS”[Title] OR “ITS1”[Title] OR “ITS2”[Title])’

else:

> gene_query = f’(“{gene}”[Gene] OR “{gene}”[Title] OR “{gene}”[All Fields])’

In addition, search results must not include whole genome shotgun data as the retrieved records should be short nucleotide fragments between 100 bp and 1.5 kb. This is to ensure that the output is compatible for clustal-omega alignment for the subsequent step in the workflow that will be performed by the second sub-agent.

> org_query = f’(“{organism}”[Organism] OR “{organism}”[All Fields])’
>
> exclusions = ‘NOT wgs[Property] NOT “whole genome shotgun”[Title]’
>
> query = f’{org_query} AND {gene_query} AND 100:15000[SLEN] {exclusions}’

### Aligning short sequences in sub-agent 2: Alignment Agent

Sub-agent 2 will submit the sequences for multiple sequence alignment using online tool via API tool calling at https://www.ebi.ac.uk/Tools/services/rest/clustalo. The job will be in queued and processed on the European Bioinformatics Institute (EBI) server. Once completed, sub-agent 2 will ensure that it is compatible FASTA format. The FASTA file will be passed to sub-agent 3 and back to the user to download.

### Finding consensus region within the aligned sequences in sub-agent 3: Analyst Agent

The alignment file will be examined using a sliding window of range of 20 to 31 nucleotides. It will also remove the dashes and gaps in the aligned window. If the candidate region after gap removal is less than 18 nucleotides, the candidate sequence will be dropped. The Analyst Agent also allows degenerate sequences up to a limit of 5 N, any candidates that exceed 5 will be rejected. Too many Ns make the sequence unreliable. The agent then counts the number of sequence records that match the window using the conservation ratio: Ratio = matches/len(target_msa). Only matches with a ratio of more than 0.7 are accepted. Shortlisted conserved targets are saved with its starting alignment position and length after gap removal. The GC content is determined by the number of G and C out of the total window length. The melting temperature is calculated based on Biopython’s nearest-neighbour model.

### Bifurcation pathways leading to either qPCR or LAMP agents or both

The consensus sequence obtained from Analyst Agent is parsed into qPCR and LAMP agents. The qPCR step is handled by sub-agent 4 (Screening Agent) and 5 (Probe Agent). The Screening Agent is responsible for searching the forward and reverse primer pairs while the Probe Agent then looks at the results of the Screening Agent to find good matches between the forward and reverse primer loci as potential TaqMan probes. The task was divided into two agents to avoid excessive nesting of for loops that extend beyond two levels to reduce computational complexity and time.

Concurrently, consensus sequences are also simultaneously handled by sub-agent 6, the LAMP agent. The gap between primers needs to be strictly enforced. For each primer locus, we set a minimum and maximum window size for the 5 gaps between the 6 targeted primers, d_F3_F2_min, d_F3_F2_max = 0, 60; d_F2_F1_min, d_F2_F1_max = 25, 100; d_F1_B1_min, d_F1_B1_max = 5, 80; d_B1_B2_min, d_B1_B2_max = 25, 100; d_B2_B3_min, d_B2_B3_max = 0, 60. A nested for loop was employed to ensure that the conserved sequence meets the minimum and maximum size based on the start and end loci.

> for i in range(n):
>
> r1 = records[i]
>
> for j in range(i+1, n):
>
> r2 = records[j]
>
> if r2[‘Location’] - (r1[‘Location’] + r1[‘AlignWidth’]) > d_F3_F2_max: break
>
> if r2[‘Location’] - (r1[‘Location’] + r1[‘AlignWidth’]) < d_F3_F2_min: continue

This is repeated for each region, and the reverse complement is then computed for the 3 reverse primers. A set of six primers consists of F3, FIP, LoopF, LoopB, BIP and B3 will be returned to the user. Last but not least, results from both qPCR and LAMP agent nodes will then converged to a report agent, where the LLM either OpenAI GPT-4o or Qwen2.5-1.5B will summarise the status and overall result with the following prompt “Summarize these results and explain why their biological properties (Tm, Amp size, spatial layout) make them good candidates for experimental validation in 3-4 sentences. Do NOT include markdown blocks.”

## DISCUSSION

**Fig 2** shows a screenshot of the prompt “Design Taqman and LAMP assay for ITS in *Ganoderma*”. **Fig 3** shows another user input “Design Taqman and LAMP assay for recA gene in *Burkholderia*”. The LLM will extract the keywords based on the user request and proceed with the workflow to search for primer candidates. A list of the primer and probe sequences for the search used for testing can be found in the appendix section. LAMP assay typically requires a longer consensus sequence than TaqMan qPCR assay as a 6-primer LAMP assay requires recognition of 8 distinct target sites while a TaqMan only require three, the forward and reverse primers with an internal probe. The FIP and BIP LAMP primers are about 40-50 nucleotides long [18]. A typical LAMP assay would require about 300 bp consensus sequence while TaqMan requires less than 200 bp in length.

**Fig 2.**
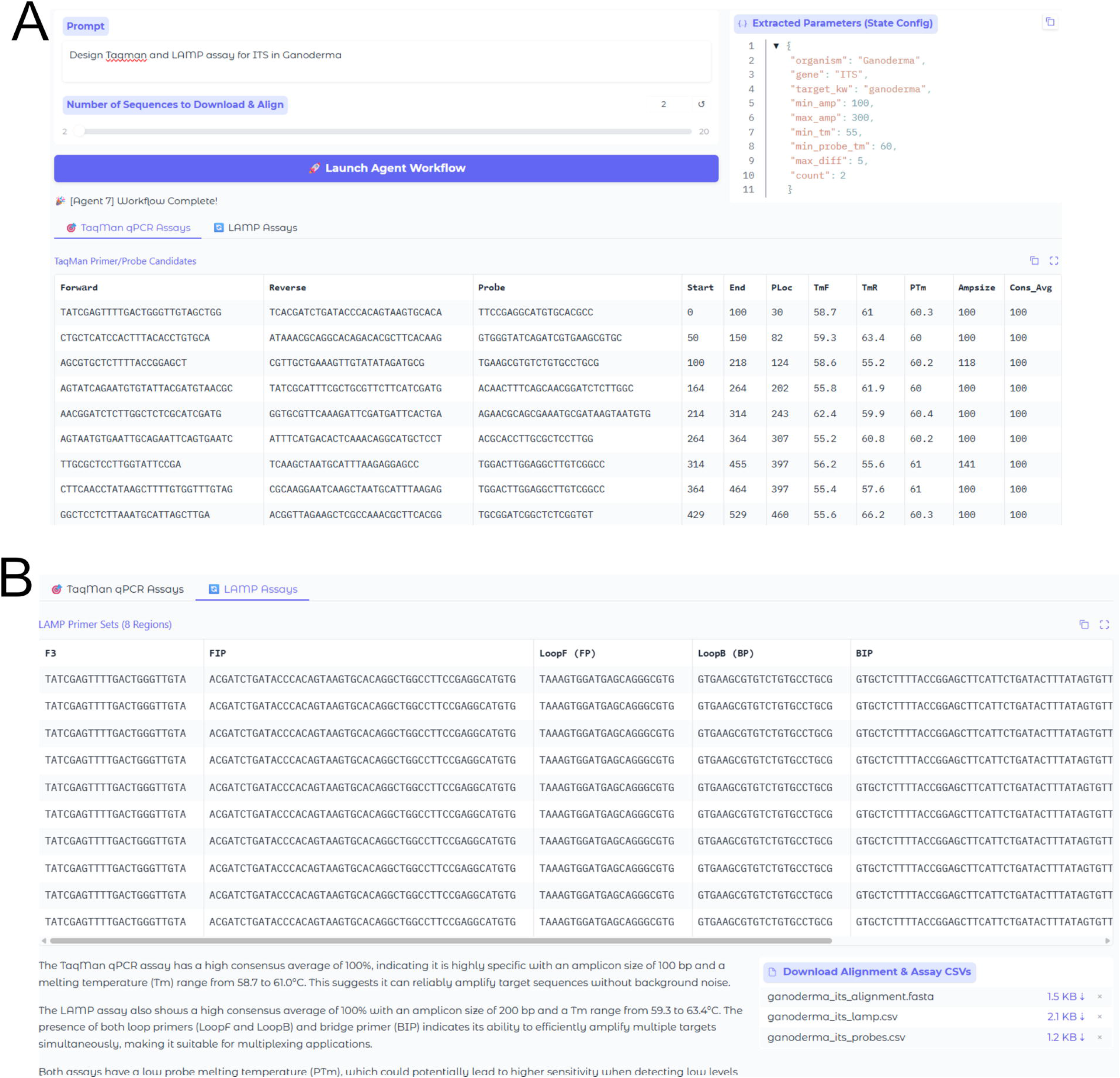
**A**. Screenshot of the user’s prompt “Design Taqman and LAMP assay for ITS in Ganoderma” and the corresponding output showing candidate primers for Taqman qPCR assay. **B**. Candidate primer sets generated for the LAMP assay. The output also provides the rationale and supporting explanations for the selection of the proposed sequences.

**Fig 3.**
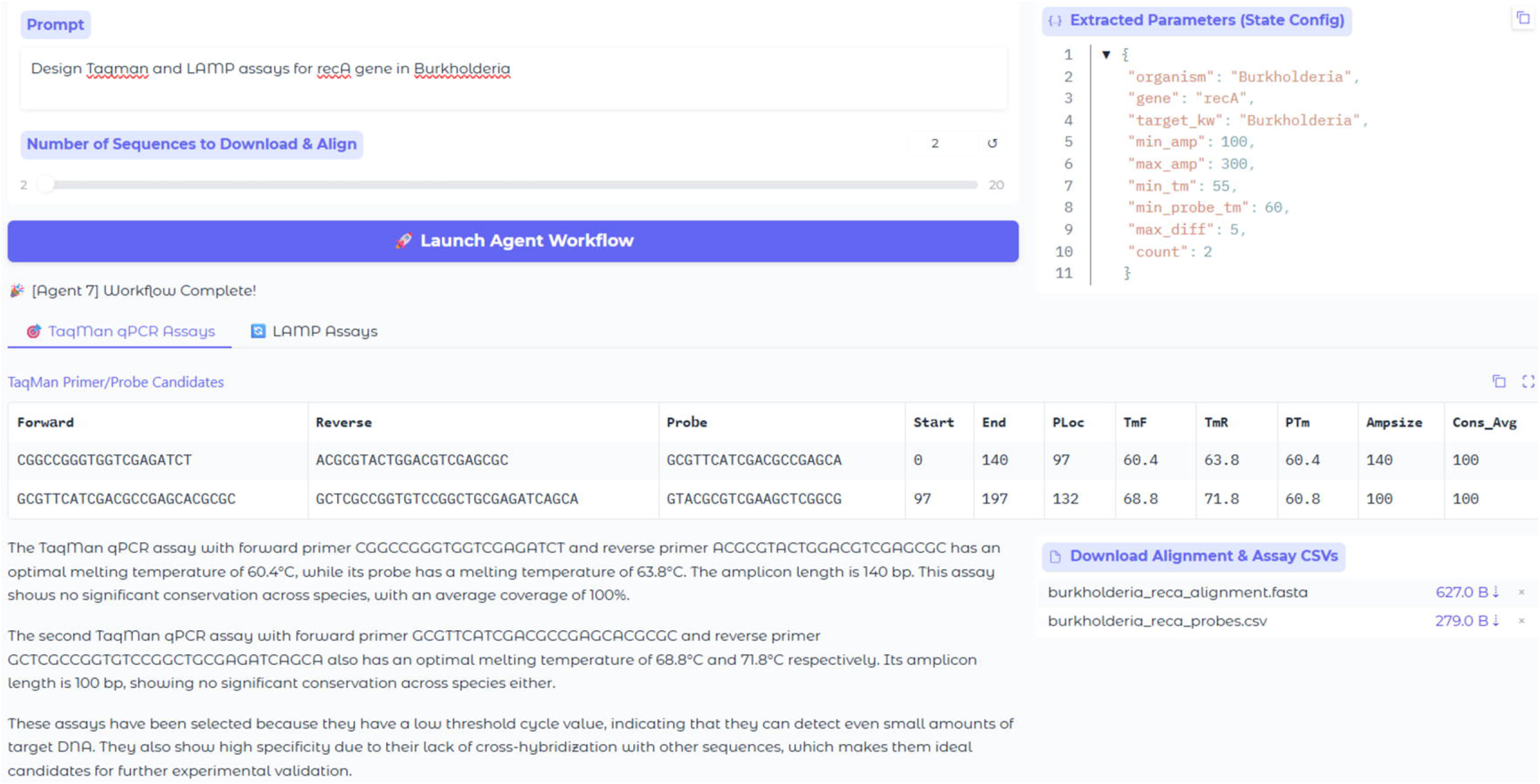
A screenshot of the user’s prompt “Design Taqman and LAMP assay for recA gene in Burkholderia”. The system identified only two qPCR assay candidates and did not generate any candidate designs for the LAMP assay.

One of the key input parameters is the ‘number of sequences to download and align’ with a default setting of 2. User can increase it by sliding the slider bar or enter the number of up to 20 sequence records. As the number of sequences to be aligned increases, the required computation time also grows considerably, depending on the user’s experimental objectives and needs. The number of sequence records to download and align was intentionally set to be variable as different search strings may yield different results for each organism in NCBI. For a well-characterized microorganism like *Escherichia coli* or *Penicillium citrinum*, a large number of sequences are readily available in NCBI databases, making sequence alignment and consensus sequence generation relatively straightforward. However, the challenge arises with less studied microorganisms like *Xylaria bambusicola*, for which NCBI may return only a very limited number of sequence records. In this case, finding the consensus sequence for at least two sequence records may sometimes be the only option available. It is advisable to refine the prompt by broadening the search to the genus taxonomic level instead of species level. The prompt in **Fig 2** where the user specifies only *Ganoderma* genera without providing the species name. In the *Burkholderia* case in **Fig 3**, it was difficult to find a consensus sequence within the aligned *recA* gene that match and fit all the criteria in the LAMP assay. Therefore, the tool only returned two sets of qPCR TaqMan primers with no LAMP primers.

## METHODS

**Fig 1** outlines the multi-agentic system workflow. The pipeline begins with user-provided inputs consisting of an organism and target gene. Keywords can be extracted using a Large Language Model (LLM). A Search Node (SearchAgent) retrieves raw nucleotide sequences, which are subsequently processed by an Align Node (AlignmentAgent) to generate a multiple-sequence alignment in FASTA format. The aligned sequences are then evaluated by an Analyze Node (AnalystAgent) to identify candidate target regions and produce a candidate dataset. Based on these candidate regions, primer design is then executed in parallel through two independent pathways: a qPCR Node integrating Screening and Probe Agents for qPCR assay development and a LAMP node (LAMPAgent) for loop-mediated isothermal amplification (LAMP) primer design. The resulting qPCR and LAMP primer sets are exported as separate csv files. Lastly, a Report Node powered by an OpenAI gpt4o-mini which consolidates the results and generates a summary. The architecture in **Fig 1** is LLM agnostic. The same agentic system was tested using Qwen2.5-1.5B instead of OpenAI GPT4o model on hugging face (https://huggingface.co/spaces/kennylau91/primerdesign). A lightweight open-source LLM is sufficient for this case and is available on a free tier.

## CONCLUSIONS

We have designed and tested our new TaqMan and LAMP software for identifying candidate primers and probes. The tool is open source and is downloadable from GitHub at https://github.com/kjxlau/primerdesign. The app is also hosted online on hugging face at https://huggingface.co/spaces/kennylau91/primerdesign. The tool aims to provide a convenient all-in-one platform for primer design, eliminating the need to download and navigate multiple standalone tools and databases. The resulting candidate primers can serve as valuable starting points for the development of point-of-care diagnostic assays.

## DATA AVAILABILITY

All sequence alignment and candidate primer sequences files for *ITS* gene in *Ganoderma, recA* gene in *Burkholderia* are available in the supplementary information.

